# COPLA, a taxonomic classifier of plasmids

**DOI:** 10.1101/2020.12.15.422809

**Authors:** Santiago Redondo-Salvo, Roger Bartomeus, Luis Vielva, Kaitlin A. Tagg, Hattie E. Webb, Raúl Fernández-López, Fernando de la Cruz

**Affiliations:** Instituto de Biomedicina y Biotecnología de Cantabria (IBBTEC), Universidad de Cantabria-CSIC, C/Albert Einstein 22, 39011 Santander, Spain; Departmento de Ingeniería de las Comunicaciones, Universidad de Cantabria, Santander, Spain; Centers for Disease Control and Prevention, 1600 Clifton Road, Atlanta, USA; WDS, Inc., 1600 Clifton Road, Atlanta, USA

## Abstract

The Plasmid Taxonomic Unit (PTU) concept is an initial step for a natural classification of plasmids. Here we present COPLA, a software for plasmid assignation to existing PTUs. To assess its performance, we used a sample of 1,000 plasmids missing from its current database. Overall, 41% of samples could be assigned an existing PTU (63% within the most abundant order, *Enterobacterales*), while 4% of samples could help to define new PTUs once COPLA database was updated.

## Introduction

Plasmids are important elements in the dissemination of genes such as antibiotic resistance determinants. The shortrange evolution of bacterial genomes (as in epidemiological outbreaks) occurs more often by acquisition of mobile genetic elements carrying, for example, resistance determinants, than by point mutations creating new alleles with a selective advantage (1). Recently we reported a procedure for the taxonomic classification of plasmids (2), and we defined PTUs (plasmid taxonomic units) as the equivalent to plasmid species. A simple and rapid method for automatic assignation of plasmids to PTUs will help in the definition of the plasmid species responsible for outbreaks of antibiotic resistant strains (3).

## Methods

### Plasmid clustering algorithm (sHSBM)

The PTU definition algorithm was based on a DNA homology network constructed from the pairwise comparison of all RefSeq84 plasmids (NCBI, dataset from September 2017), using Average Nucleotide Identity (ANI), as detailed in (2). An edge was drawn between two plasmid nodes if homology was detected over 50% of the shorter genome length. The graph-tool library was used to apply Hierarchical Stochastic Block Modeling (HSBM) to this network, with plasmids clustering based on the statistical significance of the graph topological information (4). HSBM clusters were further divided into components as, conceptually, PTU members should present DNA homology. Only clusters with at least four members were considered representative enough to be included in the next step. A home-made algorithm was added in (2) to join the previously divided clusters into a single PTU if two biological specifications were met:

1. Size compatibility: the difference on median size of both clusters is less than half of the larger median size.
2. Intercluster density: two HSBM clusters are joined if the number of edges between them is >50% of the maximum number of edges between both clusters, adjusted for their relative density.

As an example, 76% of the *Enterobacterales* plasmids in Ref-Seq84 could be assigned a PTU. More details are given in Suppl. Table 1.

### PTU prediction algorithm (COPLA)

The algorithm for predicting the PTU of new plasmids is based on the ANI homology network, processed by HSBM, and tuned as explained above. First, ANI between the query and database plasmids is calculated. Draft plasmid genomes are supported by concatenating their contigs. Next, a search for the plasmid relaxases is performed using MOBscan (5) to provide the plasmid MOB class. If not provided by the user, amino acid sequences of plasmid CDSs are predicted with Prodigal (6). Mating pair formation (MPF) and plasmid replication typing is performed using CONJscan (7) and PlasmidFinder (8). Antimicrobial resistance (AMR) genes are identified with a blastn search (>80% identity, <1e-20 e-value) against the CARD database (9).

After data input, the query plasmid is inserted into the sHSBM plasmid network described above. Next, 1,000 iterations of a multilevel Monte Carlo algorithm are performed to get a new graph partitioning that improves the Minimum Description Length (MDL) of the graph (4). The Description Length is the amount of information required to describe a graph, based on its partitioning into different clusters. It is calculated by turning the probability distribution of the different graph’s partitions into entropies. This search is constrained by impeding the creation or deletion of new partitions to ensure that the query is assigned to the already described PTUs. However, as MDL optimization is not restricted to the query but is a global process and, in addition, query introduction will change the underlying network’s topology, partition changes may be larger than expected. Finally, the HSBM output is transformed into PTUs using the same requirements described above. The query is assigned the most frequent PTU label of the members of the updated cluster. The PTU assignation is scored based on the partition overlap (10) between all database plasmids with an annotated PTU appearing in the updated query’s cluster, which indicates how much the clustering has changed due to inclusion of the query.

The COPLA output provides several files, the result of ANI comparisons, a list of plasmids related to the query and, if not provided by the user, a file with the plasmid ORFome computed by Prodigal. In addition, it provides the user with information about the plasmid relaxase, MPF type, replication formula, AMR genes and predicted PTU host-range.

## Results

We tested COPLA performance with 1,000 randomly chosen plasmid sequences uploaded to NCBI since RefSeq84 was released (see Table 1). For this we downloaded RefSeq200 release (23,309 sequences) and removed those from RefSeq84 release. Sequences with NG accession numbers were further removed as these are genomic regions, resulting in a dataset of 12,561 plasmids (Suppl. Table 2). Three possible outcomes may occur (more details in the GitHub README.md file):

1. A plasmid sequence can be assigned to an existing PTU. This happened globally in 408/1,000 cases (41%), or 259/409 (63%) for the most abundant host order (*Enterobacterales*). Output is expressed as the PTU to which the plasmid belongs, plus the PTU prediction score.
2. A plasmid sequence clusters into a group of plasmids with no previously assigned PTU. This is an interesting situation as this plasmid could be part of a potentially new PTU. Output gives the plasmids related to the query. This happened in 41 cases (4%) – or 19 cases (5%) for the *Enterobacterales*.
3. A plasmid sequence clusters with less than 4 plasmids and, so, it is below the algorithm’s minimum threshold for PTU definition.

**Table 1.**
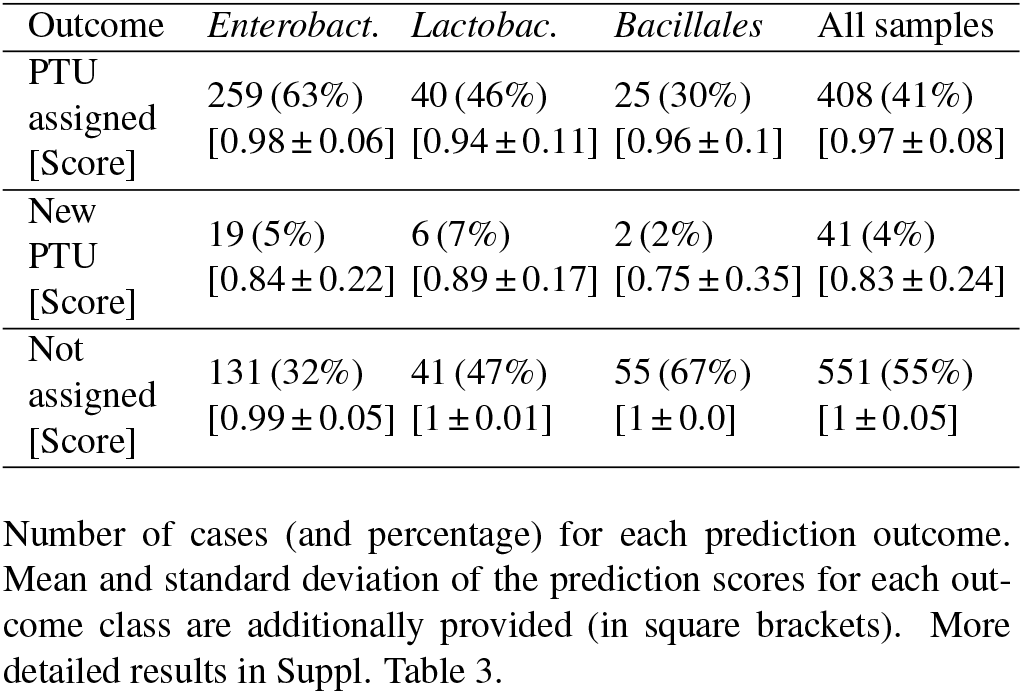
Benchmark for 1,000 new plasmids of RefSeq200 dataset for the most abundant bacterial orders

## Availability and Implementation

Source code is freely available at https://github.com/santirdnd/COPLA under the GPL v3.0 license. An online service is available at https://castillo.dicom.unican.es/copla.

## Supplementary information

Suppl_Table_1.xlsx: Metadata of RefSeq84 plasmids used for sHSBM clustering and PTUs definition.

Suppl_Table_2.xlsx: Metadata of RefSeq200 plasmids used for COPLA performance evaluation.

Suppl_Table_3.xlsx: Detailed results for the PTU assignation of the samples.

